# Identifying a ubiquitous gene expression variation pattern in the human brain

**DOI:** 10.1101/2024.07.21.604497

**Authors:** Chien-Ming Lo, Niall W. Duncan

## Abstract

The availability of dense sampled gene expression data from across the human brain has allowed important investigations into fundamental brain principles. Correlated expression patterns of genes tied to microstructural properties have been studied for their relationships with diverse brain features. This work looks at the specificity of these relationships based on the sets of genes targeted. We find that the same spatial pattern emerges from any set with more than 180 genes in it. Looking at the association between this pattern and cortical myelination, as represented by a T1w/T2w map, we show that correlations between myelination and theoretically guided gene sets do not differ from random ones. This observation prompts a reevaluation of current methodologies and assumptions in the study of gene-brain associations. Additionally, our research highlights covariance characteristics within specific functional networks, underscoring their significant role in shaping the spatial transcriptomic patterns in the brain. These findings contribute insights into the complex relationships governing brain organization and gene expression patterns. They also highlight an important methodological factor influencing the validity of inferences made from gene expression covariance patterns.

## Introduction

Understanding the relationship between brain structure and function has been a major question for neuroscience research. Methodological advances have allowed this question to be investigated in the human brain in increasing detail. One such advance has been the increasing availability of localised gene expression data. Such data can allow questions to be asked about local molecular properties of brain tissue and for these properties to be compared across different brain areas or different study populations (7).

An important resource driving such research into gene expression in the human brain has been the Allen Human Brain Atlas (AHBA). This is an openly available set of mRNA expression values for over 16,000 genes, densely sampled across the whole brain (10, 11). Some prior work using the AHBA has focused on the expression of individual genes but an influential alternative approach has been to look at overall expression patterns or at patterns of relatively large subsets of genes related to some process of interest. In such analyses, low-dimensional representations of expression similarity patterns for a set of genes are calculated through techniques such as principle component analysis (PCA) or diffusion embedding. These low-dimensional representations can then be analysed further and correlated with other aspects of the brain, such as local macrostructure or functional properties.

Interestingly, analyses that focus on low-dimensional representations of different sets of genes often report similar spacial patterns. Specifically, they report a general anterior-posterior distribution, within which unimodal regions appear distinct (13). This pattern has been seen when analysing, for example, sets of genes related to neurons, oligodendrocytes, and synaptic processes (4). It is also seen when analysing all available genes in the AHBA together or all genes with elevated expression within the brain (4, 11, 31).

A similar pattern being found across different types of genes raises both methodological and biological questions. Methodologically, we can ask how this similarity in spatial distributions across gene-sets relates to the specificity of the spatial patterns obtained. In other words, if we see consistent patterns regardless of the genes being analysed, can we say that the resulting patterns represent something specific about those genes? If not, our interpretation of the pattern should not be one based on the properties of those genes in particular as the pattern could equally represent some other property inherent to both the genes analysed and those that were not. A finding of shared spatial patterns across disparate gene-sets would lead to a set of biological questions as to what it is that is shared by different sets of genes such that consistent spatial patterns emerge across them.

To investigate these questions we used the AHBA data to systematically investigate the spatial pattern of low-dimensional representations from different sets of genes. Starting with feature-specific gene-sets reported in previous literature, we then proceeded to look at the patterns emerging from entirely random collections of genes. Finding that these patterns match, we then conducted a number of analyses to better typify the spatial patterns identified. From these analyses, and with reference to prior work (13, 25), we make some tentative suggestions as to why a consistent pattern of gene expression variation is observed irrespective of what gene-set is chosen.

## Results

### Recurring spatial patterns across gene subsets

To begin, AHBA cortical microarray data were accessed and preprocessed with the *abagen* toolbox. Preprocessing was done following established recommendations and resulted in a set of expression data for 15,633 genes, mapped with interpolation to the Schaefer-400 atlas (2, 19). Only left hemisphere regions were retained for analysis (i.e., 200 regions), as the original samples are primarily from there.

Our first question was whether or not the particular subset of genes analysed affects the resulting low-dimensional representations. To investigate this, we took a variety of gene-sets related to brain features that have been described in previous work (4) and calculated low-dimensional representations of each. These were then compared with a low-dimensional representation of a set of genes not related to brain-specifc features. The brain-related gene-sets were one of genes preferentially expressed in: all usable genes(15,633 genes); the brain as a whole (1,899 genes); neurons (2,313 genes); oligodendroglia (1,610 genes); and synapses (1,738 genes). The set of genes not specific to the brain were all those remaining after excluding these brain-related ones (10,179 genes).

For each gene-set, regional expression covariance matrices were calculated and the first principal components (PC1) of these taken as low-dimensional representations of cortical expression patterns. These cortical expression maps revealed a consistent pattern across all of the gene-sets included (Figure 1). Consistent with prior literature (4, 11, 19, 20, 30, 31), sensory areas, such as the somatosensory and visual cortices, were seen to be distinct from association areas. The expression patterns seen were robust to changes in the type of expression similarity matrices used, the specific cortical parcellation, and the dimensionality reduction algorithm employed (Sup. Figure 1).

**Figure 1.**
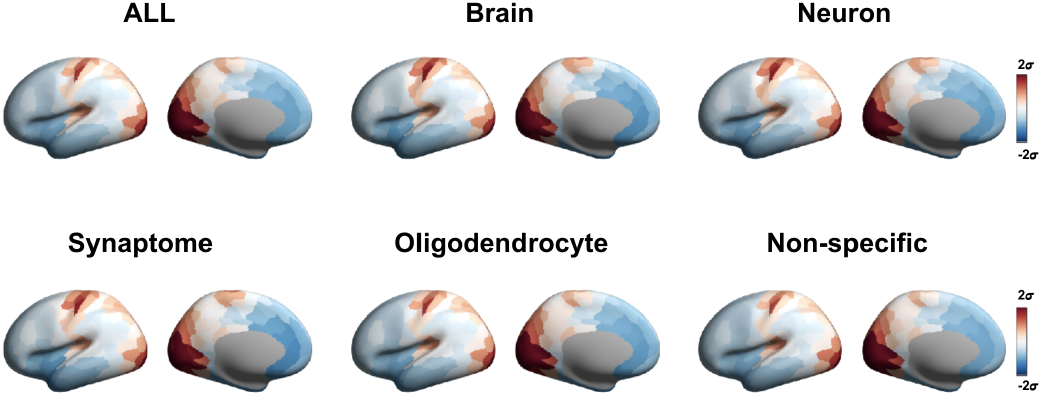
The first Principal Component of Regional Transcriptomic Similarity Across Different Gene-Sets. This figure illustrates the first Principal Components (PC) representing regional transcriptomic similarity indexed by covariance matrix across six different gene-sets on Schaefer 400 parcelation scheme. The first PC, which explains the majority of the variance (mean: 59%), appears to exhibit a universal spatial pattern regardless of the specific gene-sets. All values are standardized (z-scored) and color-coded with limits set to two standard deviations (*σ*).

The first principal component shows a general anterior-posterior distribution, within which unimodal regions appear distinct (20).

The degree of similarity between the different gene-sets was high, with the lowest correlation coefficient between brain-related sets being 0.99 (Spearman’s rank correlation). Notably, the similarity of brain-related sets to each other was not substantially different from the similarity of these with the non-brain specific gene-set (lowest *ρ* = 0.98, Spearman’s rank correlation; see Sup. Figure 2A for all correlation coefficients). This means that near-identical expression patterns are seen irrespective of what subset of genes are used an an input. The similarity between gene-sets was lower and more variant for the second (*ρ* = 0.96) and third (*ρ* = 0.89) principle components in all cases (Sup. Figure 2B).

### Establish Null Gene Distribution

The apparent lack of specificity of cortical expression patterns observed raises questions about previous findings linking the expression of particular gene-sets to other brain features. For example, the expression of particular sets of brain-related genes has been reported to correlate with patterns of myelination, as quantified through the ratio of T1-weighted to T2-weighted MR images (4, 13). If this relationship is indeed specific to particular, functionally informed, sets of genes then we would expect it to be different from that seen with randomly selected genes.

To test this, we first correlated PC1 of the brain-specific gene-set with the T1w/T2w map, obtaining an *R*^2^ of 0.69 (Figure 2A, B). This correlation was then compared to a null correlation coefficient distribution obtained by repeating this process with 5000 gene-sets of the same size (1,899) randomly selected from the non-specific gene-set. The mean of this null correlation distribution was *R*^2^ = 0.68 (95%CI: 0.66-0.69), with the empirical correlation coefficient ranking within the distribution at a percentile of 97.5 (Figure 2C). This means that the correlation of brain-specific genes with the T1w/T2w map was not different from that obtained with randomly selects non-brain genes (*α* = 0.05, two-tailed). This result was robust to the type of gene expression similarity matrix (Sup. Figure 3) and cortical parcellation used (Sup. Figure 5), although a difference between empirical and null *R*^2^ values was seen when using diffusion embedding rather than PCA as the dimensionality reduction technique (difference in *R*^2^ = 0.03, *P* = 0.042; Sup. Figure 7).

**Figure 2.**
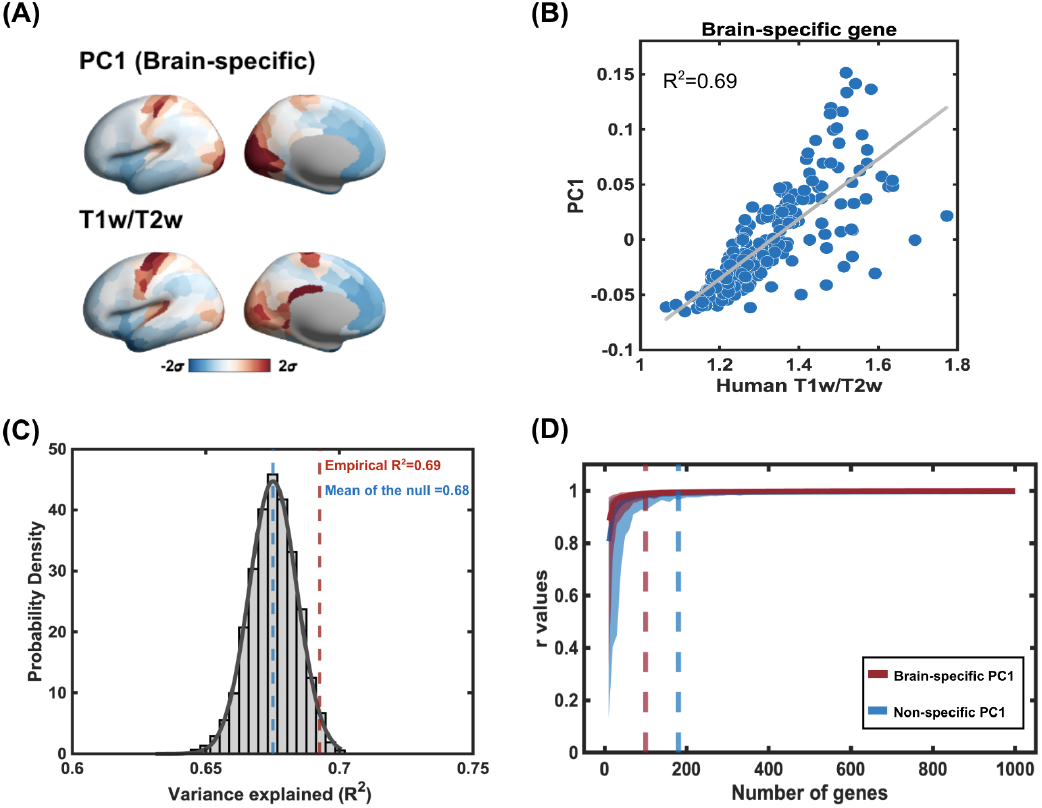
Re-evaluating the Relationship Between T1w/T2w Map and Dominant Transcriptomic Spatial Pattern Using Null Gene Distribution. **(A)**It demonstrates the replicable spatial pattern of PC1 derived from the brain-specific gene-set, akin to the human T1w/T2w map. Both maps are standardized in *σ* units. **(B)** The scatter plot on the right illustrates a strong correlation between these two maps (*R*^2^=0.69, Spearman rank correlation) as previous studies shown. **(C)** We establish a null distribution by correlating the human T1w/T2w map with PC1 derived from 5000 permutations of 1899 genes sampled from all gene-sets. The mean R2 of the null distribution is 0.68 (blue dashed line), while the empirical *R*^2^ is 0.69 (red dashed line), ranking at 97.5%. **(D)** This figure explores the emergence of the dominant spatial pattern across two gene-sets, specifically Brain-specific (in red) and Non-specific (in blue). The x-axis represents the number of genes included in re-calculating PC1 (ranging from 10 to 1000 in intervals of 10). Each gene subset is bootstrapped 500 times to estimate a 95% confidence interval (CI). The red solid line represents the average correlation values (r) between the Brain-specific PC1 and subsets of PC1, with the shaded line indicating the 95% CI. The dashed vertical line indicates the point at which an r value of 0.99 is achieved (N=100 for Brain-specific and N=180 for Non-specific).

### Emergence of the Dominant Spatial Pattern

The consistent spatial patterns observed when randomly drawing sets of genes suggests that this may be a general feature that emerges when the similarity of expression patterns of a sufficient number of genes is calculated. To investigate this supposition we systematically subsampled the brain-specific gene-set, randomly drawing from 10 to 1,000 genes at intervals of 10 (500 samples each). The first principal component of each sample’s similarity matrix was then correlated with that of the full set of 1,899 brain-specific genes. The correlation values (*r*) were first Z-transformed, averaged, and then transformed back. This analysis showed that expression patterns had a near-perfect match (mean *r* = 0.99, 95% CI: 0.97 – 1) with the full set once the sample included 100 genes or greater (Figure 2D). Randomly sampling from the set of non-brain-specific genes instead gave a similar outcome, although somewhat more genes needed to be included, at 180, to achieve the same similarity level (mean *r* = 0.99, 95% CI: 0.98 – 1; Figure 2D). All results were robust to methodological variations (see Sup. Figures 3, 5, and 7).

### The distinct covariance patterns in the functional networks

With the same low-dimensional representation of gene expression similarity arising from any sufficiently large sampling of genes, the question arises as to what the underlying brain features causing this may be. To begin to investigate this, we looked more at the properties of the similarity matrices and resulting low-dimensional representations arising from them.

Taking the expression covariance matrix, it can be seen that the Visual, Somato-motor, and Limbic networks all tend to show high covariance values (Figure 3A). Categorical regression then shows that grouping the matrix by canonical functional networks explains 53.6% (adjusted *R*^2^) of the first principal component variance (32). This analysis places the Visual (*β* = 0.08, *P <* 0.001) and Somatosensory (*β* = 0.05, *P <* 0.001) networks as being distinct at one end of the principal component and the Limbic network as distinct at the other (*β* = −0.03, *P <* 0.001; Figure 3B). This suggests that the spatial pattern of transcriptional similarity exhibits variations influenced by the intrinsic characteristics of functional networks and that these three networks are in some way distinct from the others. It is notable, though, that whilst the Limbic network has similarly high covariance values as the Visual and Somato-motor ones, it appears at the opposite end of the first principle component.

**Figure 3.**
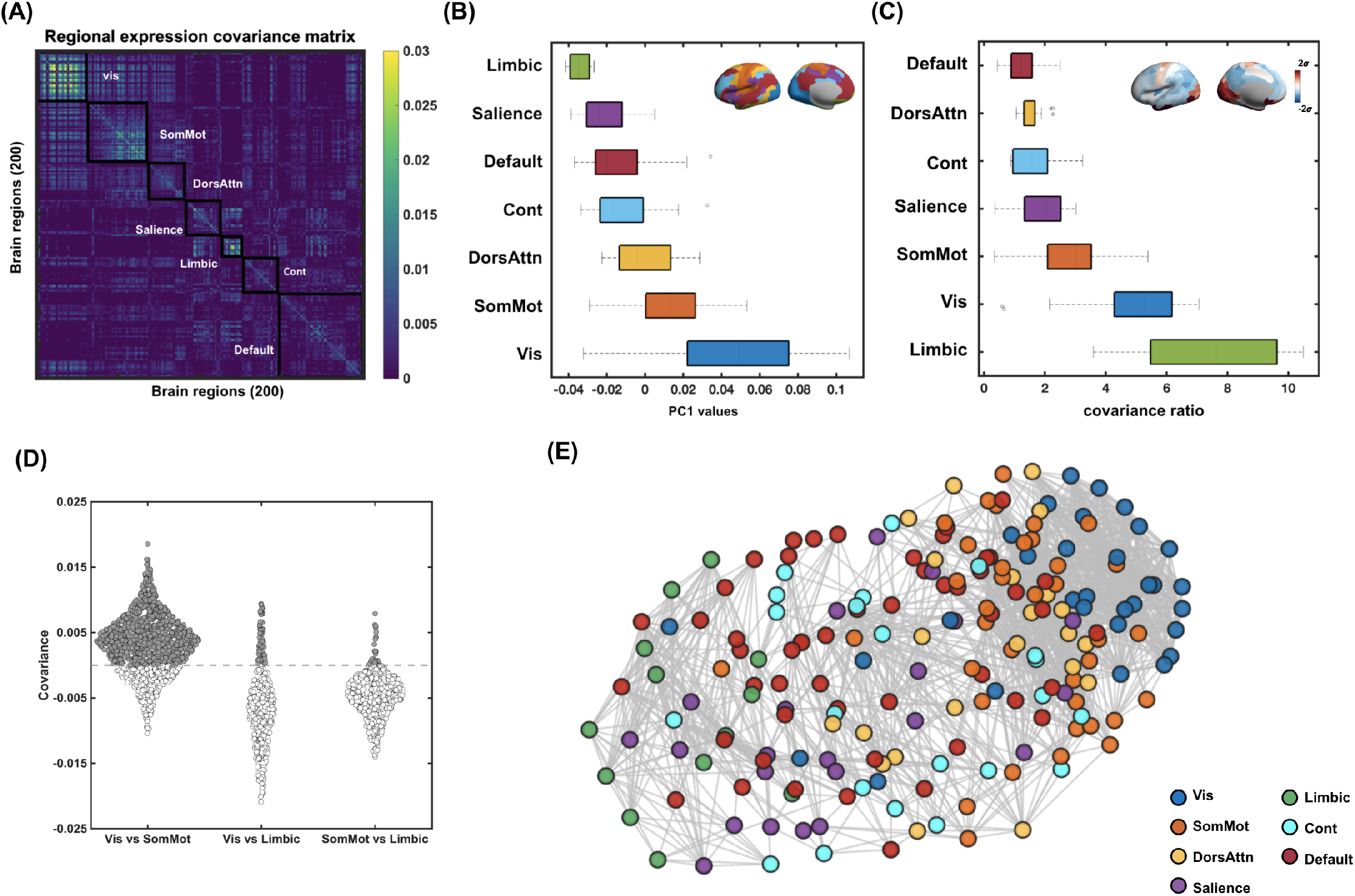
Exploring Transcriptomic Covariance Patterns Across Functional Networks. **(A)** The regional transcriptomic covariance matrix outlined by Yeo-7 functional network boundaries, with acronyms provided for reference. Key networks include Vis (Visual), SomMot (Somatomotor), DorsAttn (Dorsal Attention), Salience (Salience and Ventral Attention), Cont (Control), Limbic (Limbic), and Default (Default Mode). **(B)** Boxplot representation of PC1 values, organized and sorted by functional networks. The distribution highlights that Vis and SomMot networks are at one end of the axis, while the Limbic network is at the other. Cortical map of network assignments are color-coded in the top-right corner of the plot. **(C)** Boxplot illustrating the inter-/intra-covariance ratio, sorted by functional networks. Vis, SomMot, and Limbic networks exhibit significantly higher intra-network transcriptomic similarity compared to inter-network relationships (F(6, 65.32) = 46.05, p < 0.001). The index is visualized on the cortical surface and displayed in standard deviation (*σ*) units in the top-right corner of the plot. **(D)** The inter-network covariance comparison between Vis, SomMot, and Limbic networks. Values above zero are presented in gray. Notably, the Limbic network exhibits lower inter-network covariances between both Vis and SomMot networks. **(E)** A force-directed graph representation of the transcriptomic covariance matrix (top 90%) using the compound spring embedder (CoSE) algorithm in Cytoscape (public available at https://cytoscape.org/download.html). This visualization method reveals network relationships within the expression similarity. Notably, it reinforces the close association between the Vis and SomMot networks, while also illustrating that the Limbic network appears relatively distant, despite its high covariance pattern.

To further investigate this observation, we calculated the ratio of intra-network compared to inter-network covariance for each network. A higher index value of this would signify that a network’s transcriptomic similarity is more pronounced within itself compared to its similarity to other networks. It was found that the covariance ratio differed between networks (*F*_(6,65.32)_ = 46.05, *P <* 0.001, Welch’s ANOVA), with the Limbic network having the highest covariance ratio (7.36), followed by the Visual (5.00) and Somatomotor (2.85; Figure 3C). Post-hoc tests showed that these three networks had higher covariance ratios than the remaining four, with those four not differing from each other. Where the differ is in the Visual and Somato-motor networks having high cross-covariance with each other whereas the Limbic varies negatively with these (Figure 3D). This may explain the Limbic network’s location in the first principle component. We represented the covariance matrix using force-directed graph (Figure 3E). In this visualization, the Vis and SomMot networks clustered closely to each other at one end, whereas the Limbic network was repelled remotely due to its less similar with other networks.

Finally, we integrate the observed dominant spatial pattern of transcriptomic similarity within an evolutionary framework, utilizing cortical expansion maps from the Neuromaps toolbox (12, 20). The PC1 of genetic similarity shows a significant negative correlation with cortical expansion (*r* = –0.54, *P*_*spin*_ = 0.014), supporting the notion that regions of minimal expansion during evolution, particularly the unimodal areas, demonstrate a greater consistency in their expression patterns. Moreover, the primary histological gradient, derived from the microstructural profile covariance (27), reveals similar relationship with genetic PC1 (*r* = –0.55, *P*_*spin*_ = 0.023). These finding, coupled with the observation that expression covariance in unimodal and limbic networks aligns with specific cell types identified in the von Economo cytoarchitectural atlas (29), highlights the complex factors shaping the prevailing expression patterns (Figure 4).

**Figure 4.**
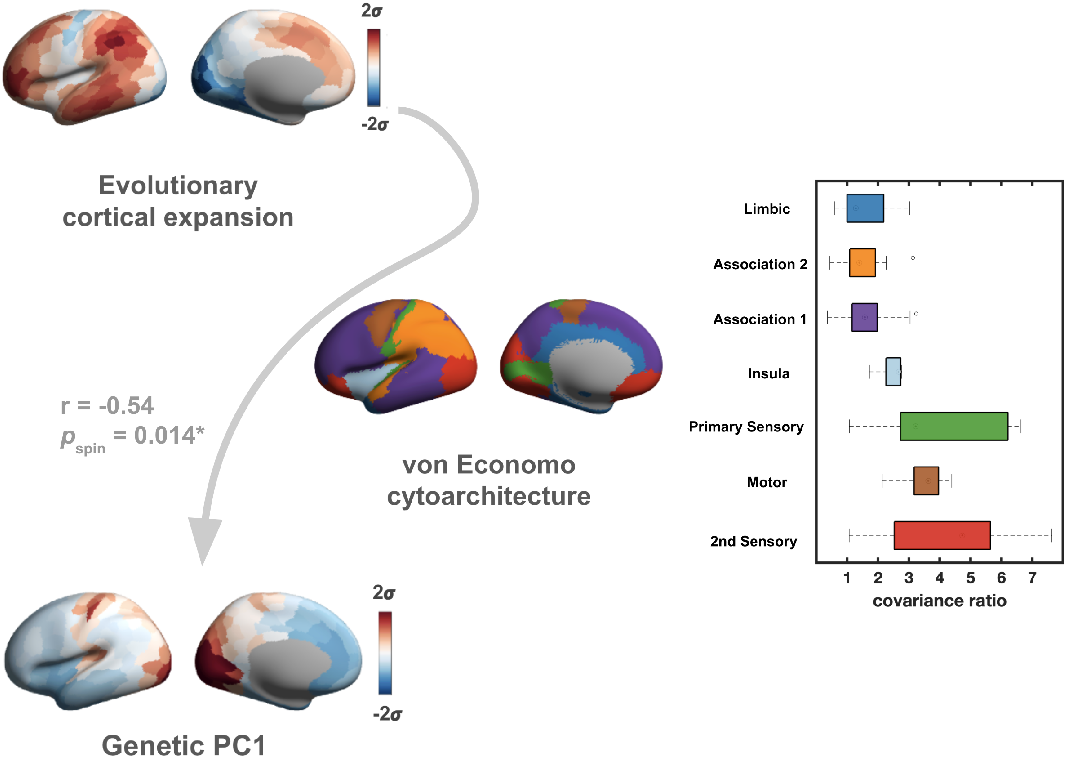
Multifaceted Influences on Genetic PC1. This figure illustrates the multifactorial influences shaping genetic PC1, demonstrating a negative correlation with evolutionary cortical expansion (*r* = –0.54, *Pspin* = 0.014), suggesting that regions with lesser expansion exhibit more homogenous expression patterns. Concurrently, the distinct network expression covariance is reflected in the von Economo cytoarchitecture, reinforcing the link between genetic profiles and cellular architecture. The convergence of genetic expression with evolutionary and cytoarchitectural data underscores the complex interplay shaping the human brain’s transcriptomic landscape.

## Discussion

These analyses show that a consistent anterior-posterior pattern emerges from low dimensional representations of expression similarity for any gene-set of sufficient size. Once a gene-set has more than around 180 different genes in it, this ubiquitous pattern appears to always be found. With the same pattern being seen irrespective of what genes are entered into an expression similarity analysis, inferences about these patterns becomes difficult. This is because the properties of the gene-set of interest cannot be immediately distinguished from a random selection. At the same time, associations between the gene-set of interest and any other measure cannot, without further investigation, be said to be due to some specific property of that gene-set, instead being a potential general feature.

These results suggest that analyses of gene expression similarity should include tests against null gene-sets of equivalent size to that being investigated. For the low dimensional representation spatial pattern itself, researchers may compare the pattern they obtain from their gene-set of interest to a set of null patterns from random gene selections. This comparison can quantify overall spatial similarity and may highlight specific areas of difference, should they exist. For association analyses between expression and other brain properties, the association strength should be tested against a null distribution produced from randomly selected gene-sets. In our own analysis, we compare the association between a canonical cortical T1w/T2w map and a brain-specific gene-set, finding that the correlation strength does not differ from a genetic null. This would suggest that there is in fact no specifically interpretable association between the target, brain-specific, gene-set and the measure of brain myelination used (in contrast to prior reports (4)).

Looking at the spatial distribution of the gene expression pattern, we observe that it follows a distinction between sensory and limbic networks compared to networks associated with more associative and integrative functions (13, 32). This concurs with prior work showing that patterns of fMRI functional connectivity, from which the network atlas used is obtained, are influenced by transcriptional similarity (17, 24). Our analysis of intra– and inter-network covariance highlighted how the sensory and limbic networks are highly self-similar in their expression patterns whereas associative networks tend to have expression patterns that are more similar across networks. This functional pattern is closely tied to structure with, for example, the limbic network having similar neural population types to sensory areas (29, 33, 35). The pattern also echos the relative degree of similarity across species for these networks, where the properties of sensory networks are more conserved than are those for associative ones (6, 17, 24, 36). This pattern of increasing expression variety in association cortex is highlighted in our work by the correlation with a map of evolutionary cortical expansion (12). Together, these factors suggest that the ubiquitous expression pattern observed stems from general patterns of cellular and organisational variation across the cortex that lead to generalised differences in measured genetic variability.

Some limitations of the work should be noted. The ubiquitous pattern identified here arises from the analysis of the AHBA, which has its own idiosyncrasies and limitations in terms of the methodology employed and the sample it is based on (11). Whether the same pattern would be seen with alternative methods or in different samples remains to be established. It may be that higher resolution sampling, taking into structural variation such as across specific cortical layers, would give a different picture (3, 14, 22). Similarly, although we ran various methodological robustness tests, it may be that novel approaches to studying large-scale gene expression similarity patterns could also produce more gene-set specific outcomes. Prior work has been criticised for incorrectly accounting for spatial proximity effects when investigating gene expression patterns (25).

To conclude, we identify a phenomenon whereby the analysis of any sufficiently large set of genes will produce highly similar spacial patterns of expression similarity. This pattern appears to reflect underlying histological variation in an non-specific manner. We suggest that any analyses of these sort should include specific tests against a genetic null to allow appropriate inferences to be made.

## Methods

### Gene expression data preprocessing

The Allen Human Brain Atlas provides regional microarray expression data from six post-mortem brains (one female and five male, ages 24–57) (10, 11). These data were processed for this work through the *abagen* and *BrainStat* toolbox (15, 19). First, microarray probes were fetched with the updated MNI152 coordinates provided by the *alleninf* package (https://github.com/chrisgorgo/alleninf). The probes were then reannotated using the data provided by Arnatkevičiūtė et al(2) and those probes that did not match a valid Entrez ID were discarded. Further probes were discarded if their expression intensity was below 50% of all samples across donors. Among probes that indexed the expression level of the same gene, the probe with the highest level of differential stability, suggesting the highly consistent pattern of regional variation across donors, was selected (10). These filtering steps left 15,633 genes that were retained for further analysis.

Following these steps, tissue sample locations were matched to regions in the Schaefer 400 parcellation (28) scheme with a distance tolerance of two standard deviations. If a brain region was not assigned a tissue sample based on the above procedure, an interpolation method was implemented using a weighted average procedure to create an estimate of the missing expression values for the region. Expression values were then normalised across samples for each gene using a scaled robust sigmoid normalisation function and rescaled to the unit interval (8). Finally, samples within each region were averaged within each donor and averaged across donors(19). To avoid sampling bias, only the 200 regions from the left hemisphere were used in subsequent analyses as only two donors have samples from the right hemisphere.

### Gene-sets

Six gene-sets were defined. The first of these consisted of all usable genes following preprocessing (15,633 genes). Four brain-related sets were then defined based on prior literature (4). These were: brain-specific genes, defined as all genes with expression levels 3SD above the median expression level (1,899); neuron-related genes (2,313); oligodendrocyte-related genes (1,610); and synapse-related genes (1,738). Note that the numbers of genes included here differ slightly from the original work as some were excluded during preprocessing (4). A final gene-set was created by removing all brain-related genes (i.e., any included in the previous four brain-related sets) from the total set of usable genes. This left 10,179 non-brain-specific genes.

### Low-dimensional representations of gene expression

A low-dimensional representation of cortical gene expression patterns was created for each of the target gene-sets from the preprocessed expression data by first calculating the covariance matrix for each set of preprocessed regional expression values. The resulting 200 × 200 matrices were then decomposed through principal component analysis (4, 11, 21, 31). The first three principal components (PC1 to PC3) for each gene-set were retained for visualisation and further analysis.

To investigate the similarity of the expression patterns resulting from each of the six gene-sets, each principal component was correlated (Pearson) with the corresponding principal component from the other sets (e.g., All genes PC1 correlated with brain-specific genes PC1, neuron genes PC1, etc.). Pattern similarity for each principal component was taken as the standard deviation across all correlation coefficients for that component.

As estimates of gene expression patterns may be sensitive to particular methodology adopted, we carried out the same analyses whilst varying key methodological elements. Firstly, we used an alternative cortical parcellation, namely the HCP-MMP1 atlas (9), to ensure that results are not overly influenced by the particular subdivision of the cortex used. Secondly, the gene similarity matrices were calculated using Pearson correlation rather than covariance as this approach is widely used in the literature. The PCA analysis was then applied to this correlation matrix. Finally, we applied diffusion mapping rather than PCA as the dimensionality reduction method. This non-linear dimensionality reduction technique is also widely used in the literature (18, 23, 26, 27, 31, 34, 37) and may capture relationships not apparent with linear techniques such as PCA. For this, the input matrix was proportionally thresholded at 90% and transformed through the application of a normalized cosine similarity kernel in order to scale the range into 0 and 1. The diffusion map algorithm was then applied to this affinity matrix to obtain a final low-dimensional embedding representation (5, 18).

### Gene-set expression pattern specificity

Particular sets of brain-related genes have been reported in prior work as being associated with structural and functional features of the brain. To investigate the dependency of such relationships upon the genes selected, we ran a spacial correlation analyses between the first PC of the brain-specific gene-set (1,899 genes) and the T1w/T2w ratio map, taken to represent myelin content, openly available at https://balsa.wustl.edu/mpwM. A null distribution of correlation coefficients between gene expression patterns and the T1w/T2w map was then calculated by randomly sub-sampling 1,899 genes from the non-brain specific gene-set, establishing the first PC of their similarity matrix, and then correlating that PC with the T1w/T2w map. This process was repeated 5000 times, giving the distribution of correlation coefficients found when random sets of genes are related to the T1w/T2w map. The correlation coefficient obtained with the brain-related gene-set was then compared to this random null distribution to test for statistical significance (*α* = 0.05, two-tailed). This process was also repeated with each of the three methodological variations described above (change in atlas, similarity estimate, and dimensionality reduction method).

### Emergence of a consistent gene expression similarity pattern

As our results suggested that highly similar gene expression similarity patterns arise regardless of which sets of genes are analysed, we investigated how many genes need to be included in a set for this consistent outcome to emerge. To do this, we first sub-sampled from the brain-specific gene-set (1,899 genes) with sample sizes ranging from 10 to 1000 (intervals of 10). The first PC for each sub-sample was then calculated and correlated with that of the full brain-specific gene-set. This process was repeated 500 times for each sample size to obtain a correlation 95% confidence interval. This process was then repeated, sampling from the non-brain-specific gene-set and relating the resulting first PCs to that of the full brain-specific gene-set.

### Features of the consistent expression pattern

Observing that highly similar gene expression similarity patterns emerge irrespective of which sets of genes are used as inputs, we carried out some additional analyses with the aim of describing features that may potentially explain this observation. In a first step, we sorted the similarity matrices and the first PC into seven functional networks (32). The relationship between this categorisation and the first PC values explained by this organisation was established through categorical linear regression. Next, we calculated a ratio of intra-network to inter-network covariance for each of the seven networks. Differences in this ratio between networks was tested through Welch’s ANOVA.

Finally, we related the gene expression pattern to underlying features of the cortex. The first of these was estimates of differential evolutionary expansion. These represent the degree to which different areas of the human cortex are thought to have increased in size through evolutionary development (12). The gene expression pattern was then related to two representations of cytoarchitectural differences across the brain, namely an atlas von Economo’s cytoarchitectural classes (29) and a histological gradient (27) calculated from the BigBrain dataset (1). The evolutionary expansion map and von Economo atlas were obtained from the *Neuromaps* toolbox (20). The histological gradient was obtained from the *ENIGMA* toolbox (16).

## Data and code availability

The analysis code for this work is available at https://osf.io/asj45/. The gene expression data used in this study were obtained from the AHBA, which can be accessed at Allen Brain Atlas. The preprocessing of the raw microarray data was carried out using the abagen toolbox and BrainStat toolbox. Additional brain imaging data and analysis tools were accessed from the ENIGMA Toolbox. The brain maps and cortical expansion data were obtained from the Neuromaps toolbox.

## Acknowledgments

This work was supported by grants to NWD from the Taiwan Ministry of Science and Technology (110-2628-H-038-001-MY4 & 113-2423-H-038-002-MY3). The preprint was created using a modified version of Ricardo Henriques’ BioRxiv LaTex template (https://henriqueslab.github.io/resources/bioRxivTemplate/).

## Contributions

CL: Conceptualization, Methodology, Formal Analysis, Visualization, Writing – original draft

NWD: Conceptualization, Methodology, Supervision, Writing – original draft

## Supplementary materials

**Supplementary Fig. 1.**
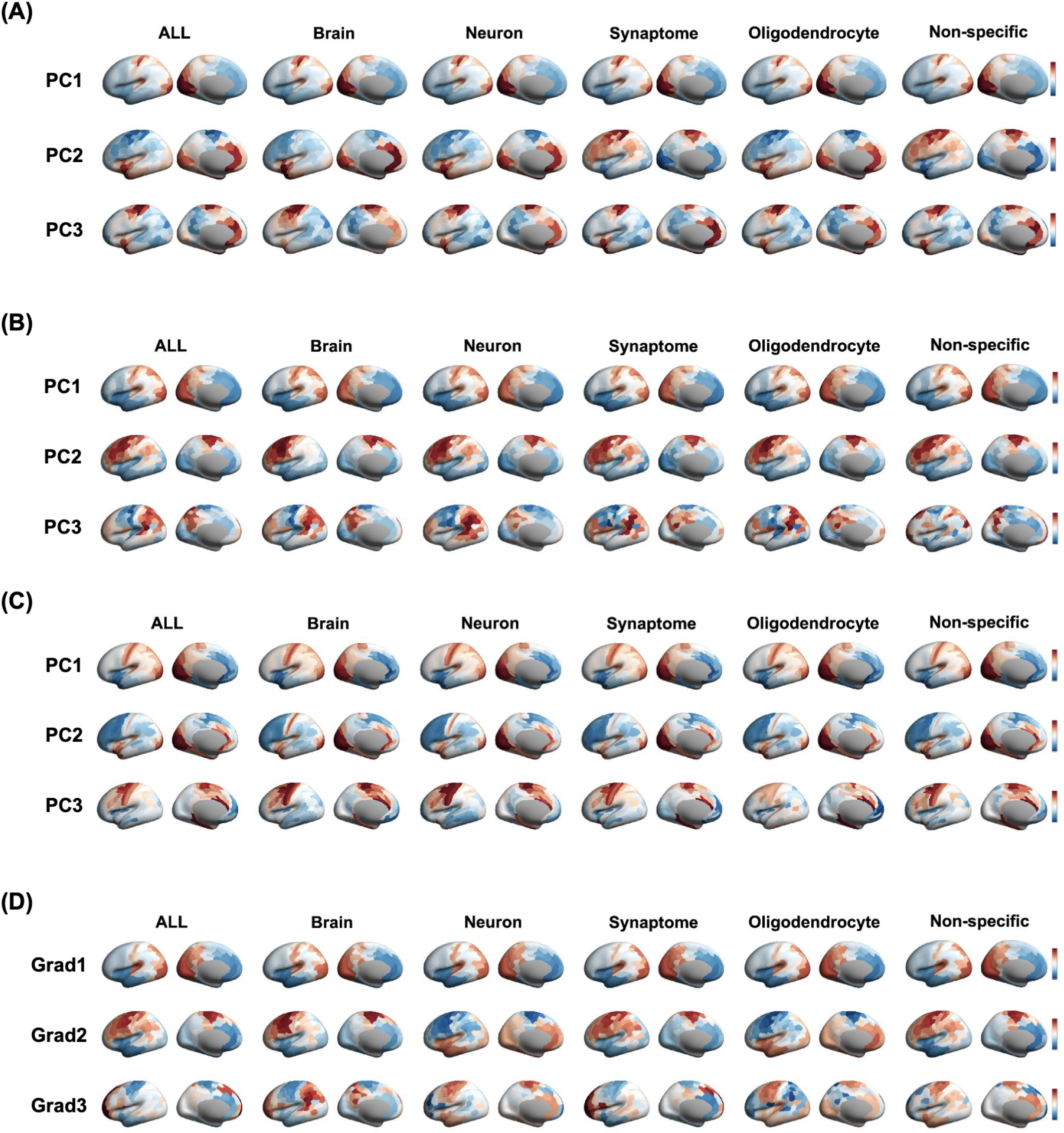
Complementary Analysis of Transcriptomic Similarity Patterns. This figure presents a series of analyses exploring the robustness of the dominant regional transcriptomic similarity patterns. Each panel showcases the first three Principal Components (PC) or Gradients derived from different data configurations. **(A)** It replicates the results presented in Figure 1, displaying the first three PCs for regional transcriptomic similarity across six gene sets using the covariance matrix on the Schaefer 400 parcelation scheme. **(B)** We maintain the Schaefer parcelation but alter the input matrix from covariance to Pearson correlation. **(C)** It maintains the use of the covariance matrix but switches to the Glasser 360 parcelation, examining the impact of parcellation scheme choice. **(D)** Finally, we explore the use of a non-linear dimensionality reduction method known as diffusion map. Here, we utilize the covariance matrix on the Schaefer parcelation with a common parameter setting, applying normalized angle and selecting the top 10% of the matrix as input. The output, referred to as the gradient (Grad), is displayed.

**Supplementary Fig. 2.**
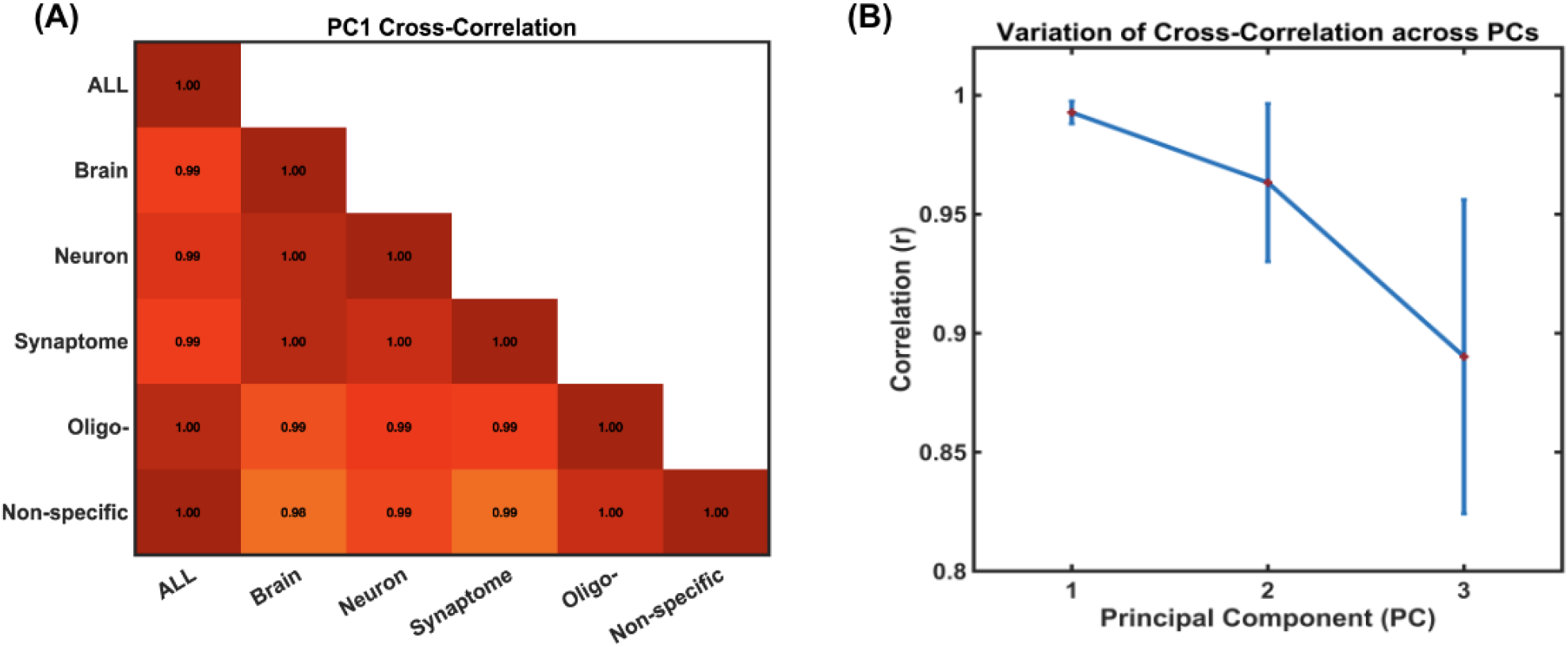
**(A)** The PC1 cross correlation between six gene-sets. The values are remarkably high with the lowest r value of 0.98 between PC1s of Brain-specific and non-specific gene-sets. The “Oligo-“ stands for Oligodendrocyte. **(B)** The errorbar plot shows the consistency of r values across first 3 PCs among different gene-sets. The pattern indicates that the variation increase as function of explained variable the PC represented.

**Supplementary Fig. 3.**
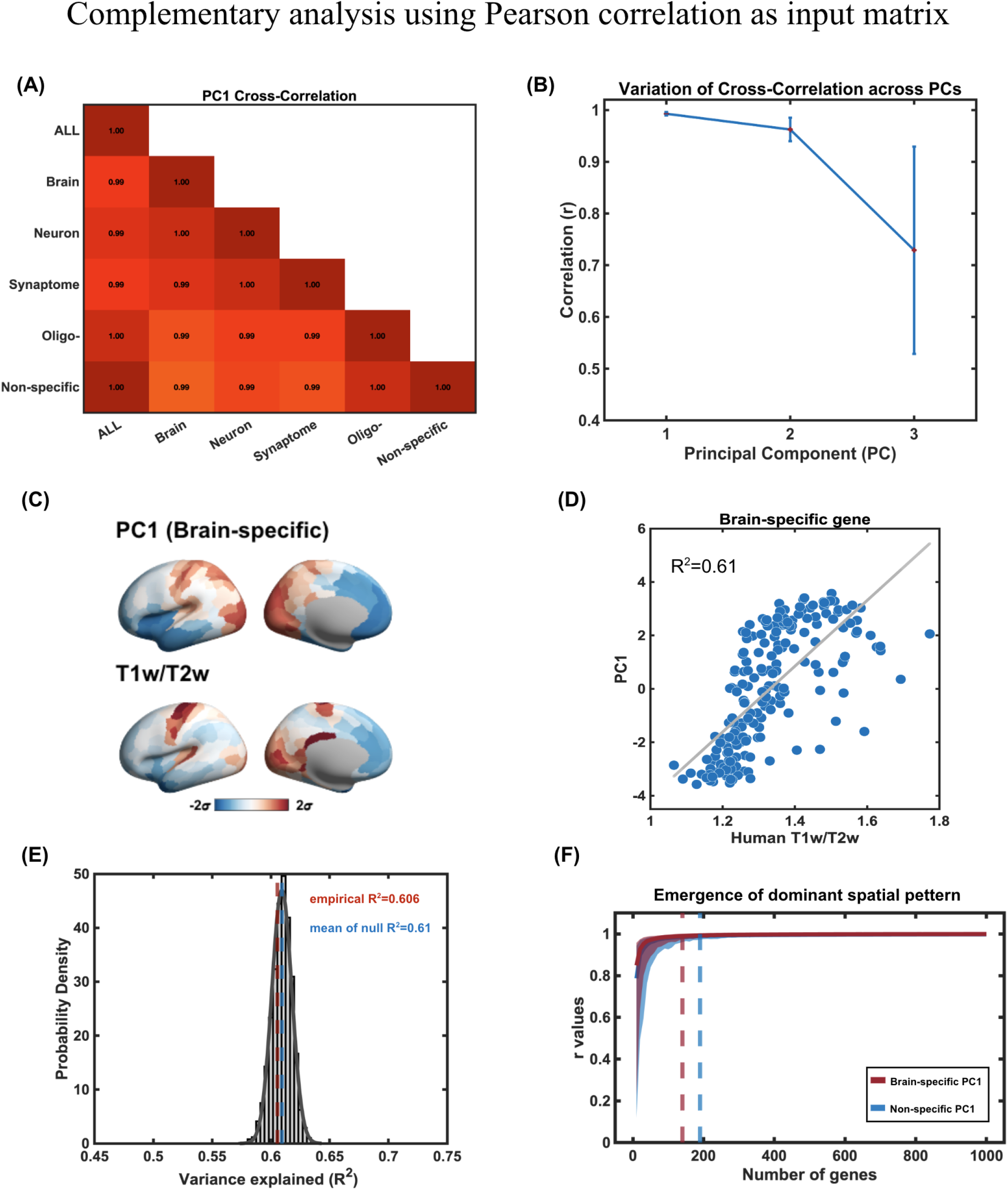
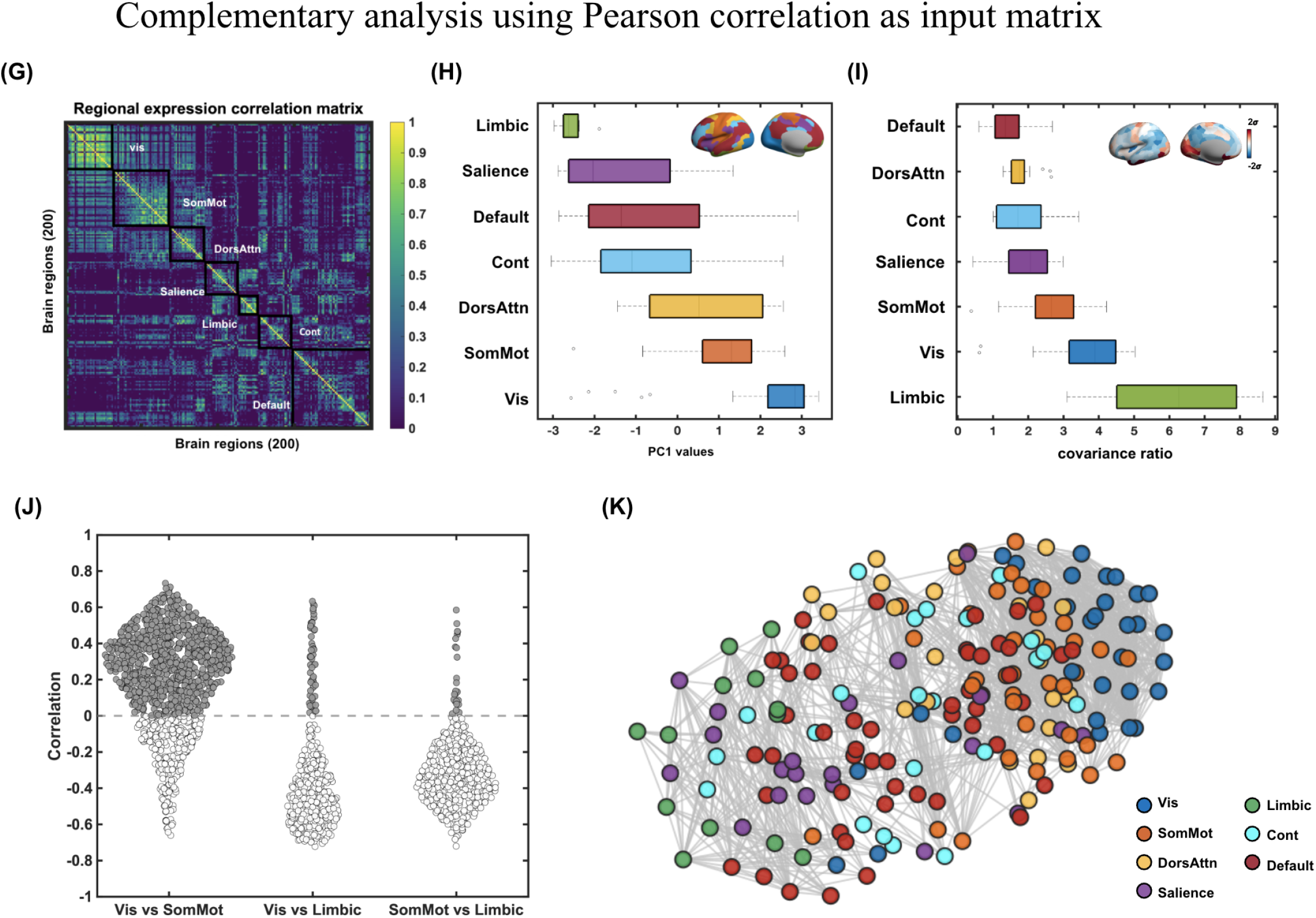
Complementary results using correlation matrix as input. **(A)** The PC1 cross correlation between six gene-sets. The values are remarkably high with the lowest r value of 0.99. **(B)** The errorbar plot shows the consistency of r values across first 3 PCs among different gene-sets. The pattern indicates that the variation increase as function of explained variable the PC represented. **(C)** The cortical maps demonstrate the replicable spatial pattern of PC1 derived from the brain-specific gene-set, akin to the human T1w/T2w map. Both maps are standardised in *σ* units. **(D)** The scatter plot on the right illustrates a strong correlation between these two maps (*R*^2^ = 0.61, *P <* 0.001, Spearman rank correlation). **(E)** We establish a null distribution by correlating the human T1w/T2w map with PC1 derived from 5000 permutations of 1899 genes sampled from all gene-sets. The mean *R*^2^ of the null distribution is 0.606 (blue dashed line), while the empirical *R*^2^ is 0.61 (red dashed line), ranking at 31%. **(E)** This figure explores whether the emergence pattern differs between gene-sets, specifically Brain-specific (in red) and Non-specific (in blue). The x-axis represents the number of genes included in re-calculating PC1 (ranging from 10 to 1000 in intervals of 10). Each gene subset is bootstrapped 500 times to estimate a 95% confidence interval (CI). The dashed vertical line indicates the point at which an r value of 0.99 is achieved (N=140 for Brain-specific and N=190 for Non-specific). **(G)** The regional transcriptomic covariance matrix outlined by Yeo-7 functional network boundaries, with acronyms provided for reference. **(H)** Boxplot representation of PC1 values, organised and sorted by functional networks. The distribution highlights that Vis and SomMot networks are at one end of the axis, while the Limbic network is at the other. Cortical map of network assignments are color-coded in the top-right corner of the plot. **(I)** Boxplot illustrating the inter-/intra-covariance ratio, sorted by functional networks. Vis, SomMot, and Limbic networks exhibit significantly higher intra-network transcriptomic similarity compared to inter-network relationships (F(6, 48.13) = 67.06, *p <* 0.001). The index is visualised on the cortical surface and displayed in standard deviation (*σ*) units in the top-right corner of the plot. **(J)** The inter-network correlation comparison between Vis, SomMot, and Limbic networks. Values above zero are presented in gray. **(K)** A force-directed graph representation of the transcriptomic correlation matrix using the compound spring embedder (CoSE) algorithm in Cytoscape.

**Supplementary Fig. 4.**
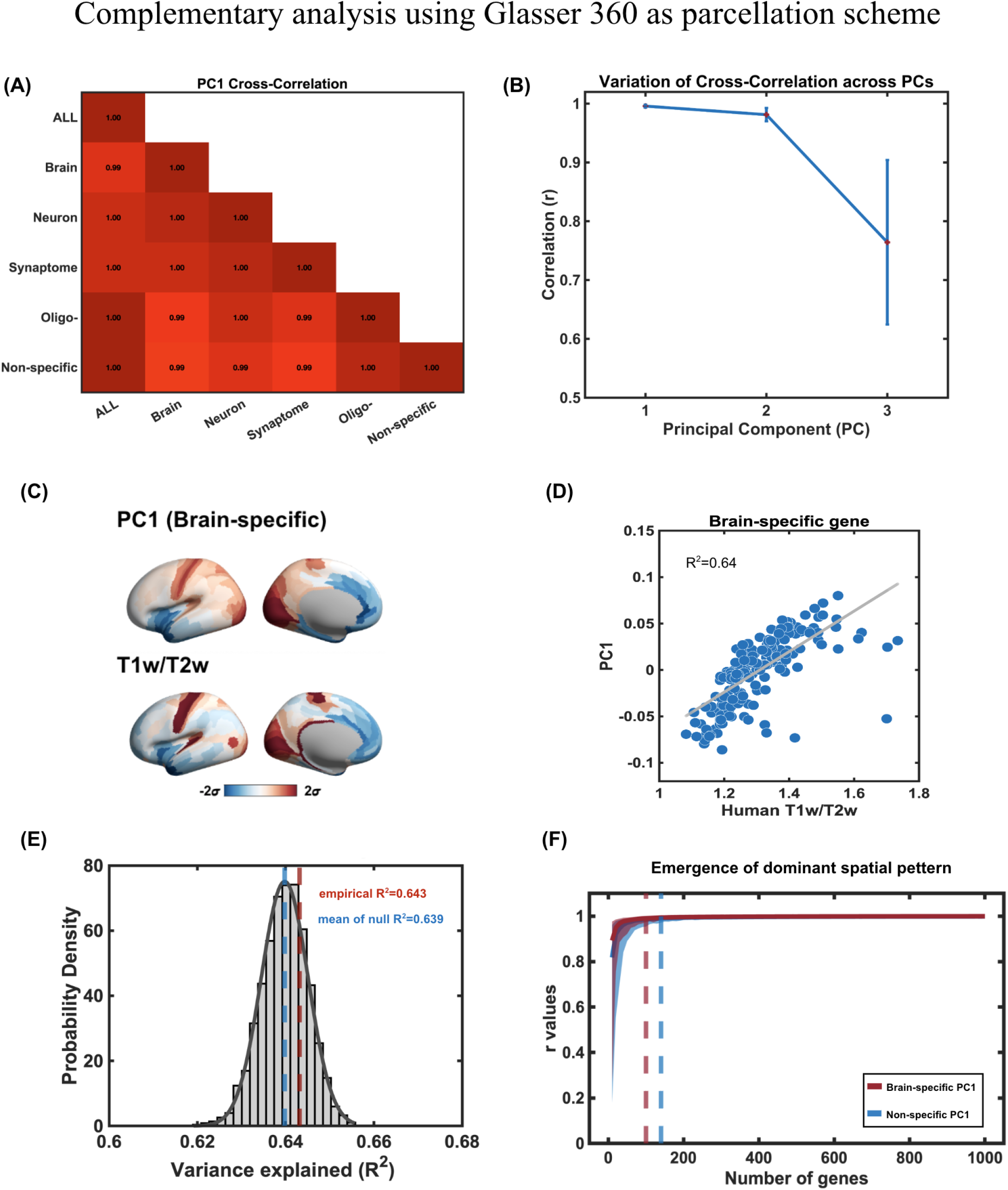
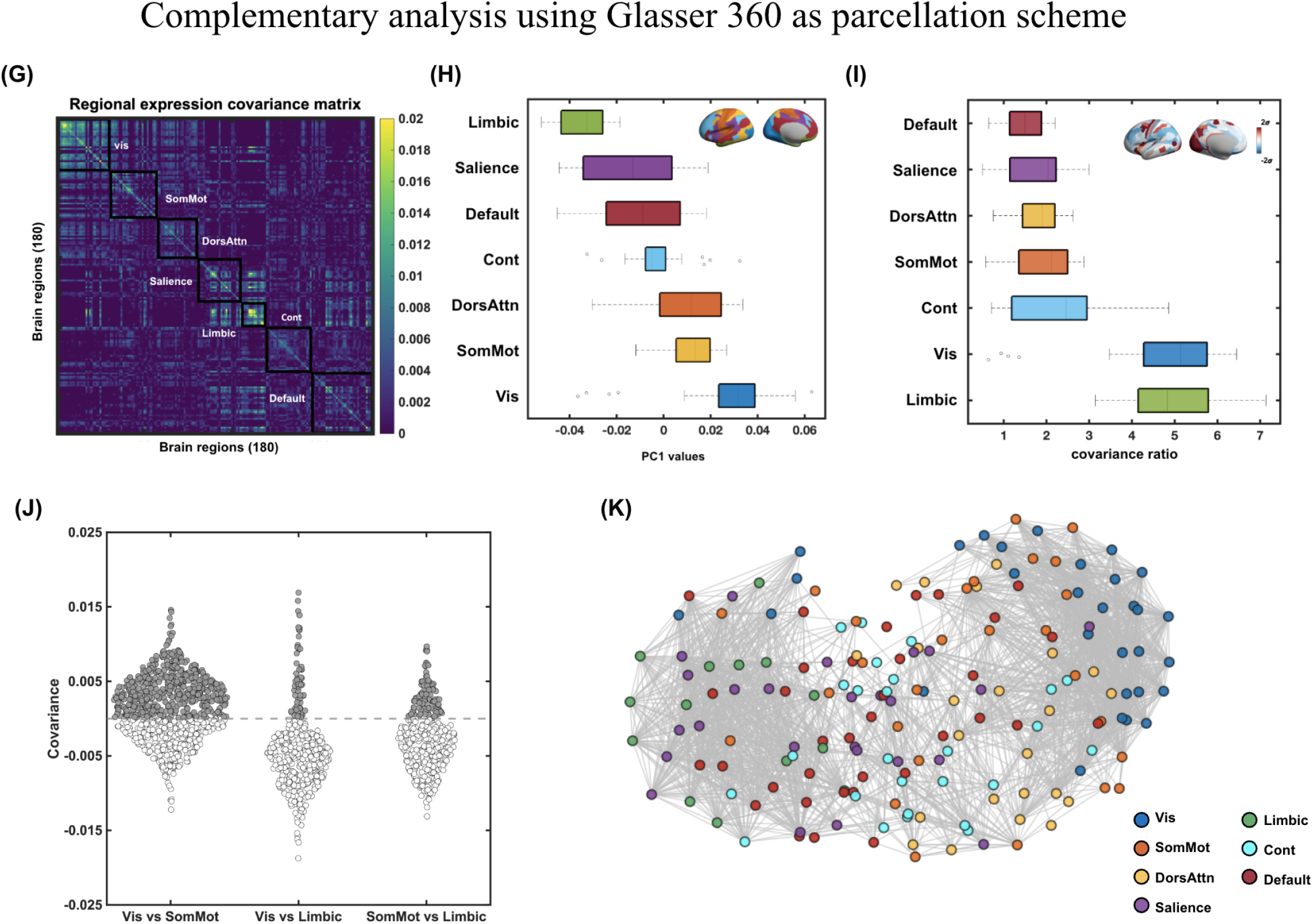
Complementary results using Glasser 360 parcellation. **(A)** The PC1 cross correlation between six gene-sets. The values are consistently high with the lowest r value of 0.99. **(B)** The errorbar plot shows the consistency of r values across first 3 PCs among different gene-sets. The pattern indicates that the variation increase as function of explained variable the PC represented. **(C)**The cortical maps demonstrate the replicable spatial pattern of PC1 derived from the brain-specific gene-set, akin to the human T1w/T2w map. Both maps are standardised in *σ* units. **(D)** The scatter plot on the right illustrates a strong correlation between these two maps (*R*^2^ = 0.64, *P <* 0.001, Spearman rank correlation). **(E)** We establish a null distribution by correlating the human T1w/T2w map with PC1 derived from 5000 permutations of 1899 genes sampled from all gene-sets. The mean *R*^2^ of the null distribution is 0.639 (blue dashed line), while the empirical *R*^2^ is 0.643(red dashed line), ranking at 73.2%. **(E)** The dashed vertical line indicates the point at which an r value of 0.99 is achieved (N=160 for Brain-specific and N=220 for Non-specific). The x-axis represents the number of genes included in re-calculating PC1 (ranging from 10 to 1000 in intervals of 10). Each gene subset is bootstrapped 500 times to estimate a 95% confidence interval (CI). The dashed vertical line indicates the point at which an r value of 0.99 is achieved (N=100 for Brain-specific and N=140 for Non-specific). **(G)** The regional transcriptomic covariance matrix outlined by Yeo-7 functional network boundaries, with acronyms provided for reference. **(H)** Boxplot representation of PC1 values, organised and sorted by functional networks. Cortical map of network assignments are color-coded in the top-right corner of the plot. **(I)** Boxplot illustrating the inter-/intra-covariance ratio, sorted by functional networks. Vis, SomMot, and Limbic networks exhibit significantly higher intra-network transcriptomic similarity compared to inter-network relationships (*F* (6, 46.38) = 49.8, *p <* 0.001). The index is visualised on the cortical surface and displayed in standard deviation (*σ*) units in the top-right corner of the plot. **(J)** The inter-network covariance comparison between Vis, SomMot, and Limbic networks. Values above zero are presented in gray. **(K)** A force-directed graph representation of the transcriptomic covariance matrix using the compound spring embedder (CoSE) algorithm in Cytoscape.

**Supplementary Fig. 5.**
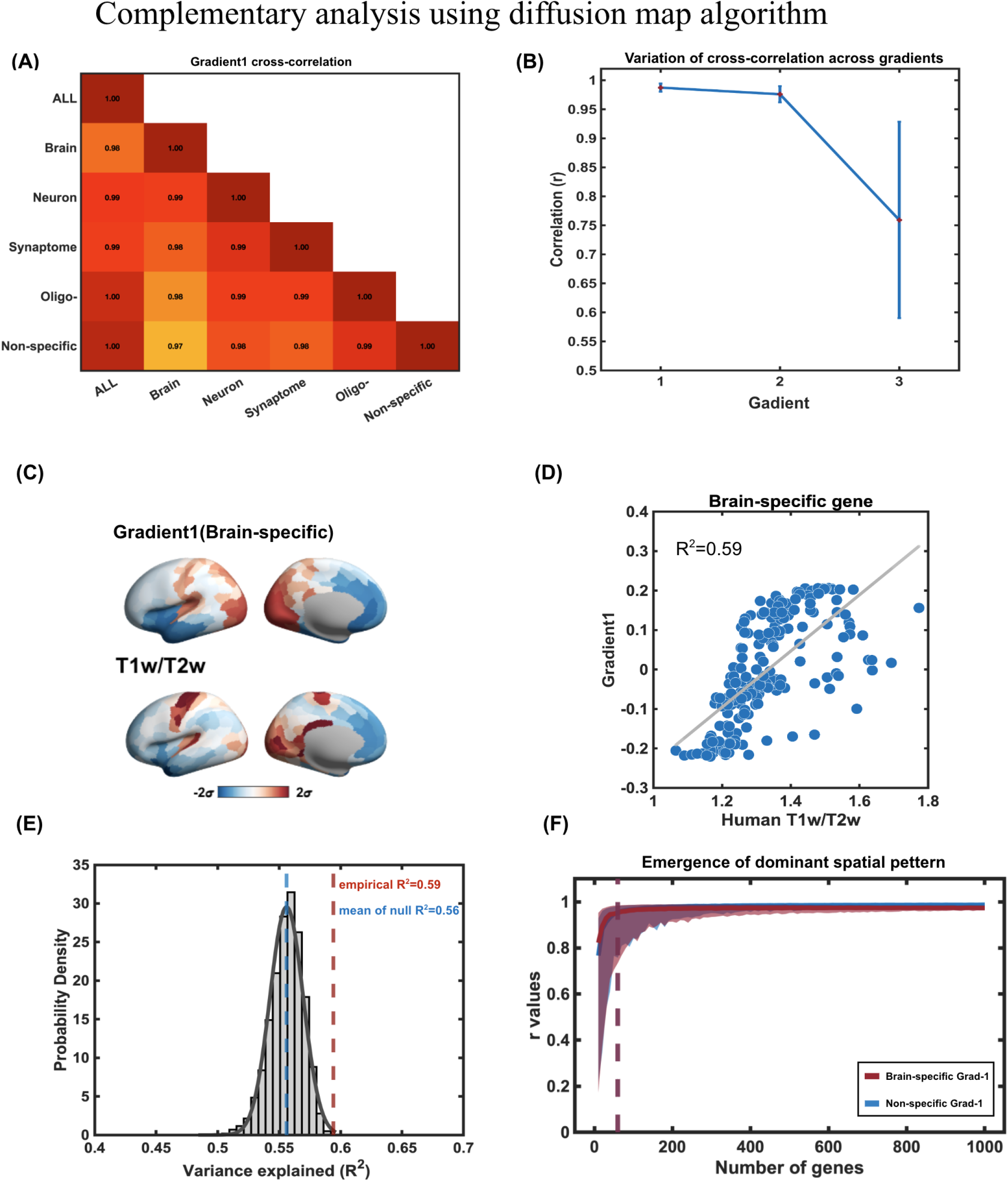
Complementary results using diffusion map method. **(A)** The Grad-1cross correlation between six gene-sets. The values are consistently high with the lowest r value of 0.97 between Brain-specific and non-specific gene-sets. **(B)** The errorbar plot shows the consistency of r values across first 3 Gradients among different gene-sets. The pattern indicates that the variation increase as function of explained variable the Gradient represented. **(C)**The cortical maps demonstrate the replicable spatial pattern of Grad-1 derived from the brain-specific gene-set, akin to the human T1w/T2w map. Both maps are standardised in *σ* units. **(D)** The scatter plot on the right illustrates a strong correlation between these two maps (*R*^2^ = 0.64, *P <* 0.001, Spearman rank correlation). **(E)** We establish a null distribution by correlating the human T1w/T2w map with Grad-1 derived from 5000 permutations of 1899 genes sampled from all gene-sets. The mean *R*^2^ of the null distribution is 0.56 (blue dashed line), while the empirical *R*^2^ is 0.59 (red dashed line), ranking at 97.9%. **(E)** This figure explores whether the emergence pattern differs between gene-sets, specifically Brain-specific (in red) and Non-specific (in blue). The x-axis represents the number of genes included in re-calculating Grad-1 (ranging from 10 to 1000 in intervals of 10). Each gene subset is bootstrapped 500 times to estimate a 95% confidence interval (CI). The dashed vertical line indicates the point at which an r value of 0.95 is achieved (N=60 for Brain-specific and N=60 for Non-specific).

